# Extensive *In Silico* Analysis of the Functional and Structural Consequences of SNPs in Human *ARX* Gene

**DOI:** 10.1101/2020.05.27.104810

**Authors:** Mujahed I. Mustafa, Naseem S. Murshed, Abdelrahman H. Abdelmoneim, Abdelrafie M. Makhawi

**Affiliations:** Department of Biotechnology, University of Bahri, Khartoum, Sudan; Department of Microbiology, International University of Africa, Khartoum, Sudan; Faculty of Medicine, Alneelain University, Khartoum, Sudan

**Keywords:** *ARX* gene, Early infantile epileptic encephalopathy 1 (EIEE1), neurologic disease, *in silico* analysis, SNPs

## Abstract

Early infantile epileptic encephalopathy 1 (EIEE1) is a rare but devastating neurologic disorder that displays concomitant cognitive and motor impairment, and is often presented in the first months of life with severe intellectual disability. The objective of this study is to classify the most deleterious nsSNPs in *ARX* gene that may cause EIEE1 disease. Despite the reported association of *ARX* gene mutations with vulnerability to several neurologic condition, there is lack of in silico analysis on the functional and structural impacts of single nucleotide polymorphisms (SNPs) of the *ARX* at protein level. Therefore, the pathogenic nsSNPs in the human *ARX* obtained from NCBI were analyzed for their functional and structural impact using bioinformatics tools like SIFT, Polyphen, PROVEAN, I-Mutant, and MUPro. The effects of the mutations on tertiary structure of the human ARX protein were predicted using RaptorX and visualized by UCSF Chimera while STRING was used to investigate its protein–protein interaction. Our extensive *in silico* analysis revealed 11 mutations that will significantly alter the structure of human ARX protein; that may disturb the domain which will affect the function of the protein. Extensive *in silico* analysis of the functional and structural consequences of SNPs in human ARX gene revealed 11 mutations (L535Q, R528S, R380L, V374D, L343Q, T333N, T333S, R332H, R330H, G34R and L33P) that may cause EIEE1.Therefore, can be used as diagnostic markers for EIEE1.

## 1. Introduction

Early infantile epileptic encephalopathy 1 (EIEE1) is a rare neurologic condition associated with both cognitive and motor impairment, and is often complicated with severe intellectual disability in neonates [1]. Patients with Aristaless-related homeobox (*ARX*) mutations have wide range of characteristics, such as agenesis of the corpus callosum, abnormal genitalia (Proud syndrome), lissencephaly, intellectual disability, and infantile spasms [2-5].

*ARX* is a paired-type homeobox gene located on the X chromosome [6]. The ARX protein is critical for the development of forebrain, pancreas and testes, and the proliferation and differentiation of cerebral interneurons [7, 8]. The most frequent *ARX* variants that cause EIEE1 are a result of polyalanine tract expansions [6, 9]. One of the complication of *ARX* gene mutation is neuroblastoma [10], which is an embryonal malignancy that dusturbs normal development of the adrenal medulla and paravertebral sympathetic ganglia in early childhood, also any shortage of ARX function is believed to affect normal brain development, leading to seizures and intellectual disability [11, 12].

Translational bioinformatics is a critical contributor in the field of personalized medicine, which targets to diminsh the gap between academic and clinical research by classifying the most deleterious nsSNPs for larger follow-up clinical studies [13-15]. In this work, the outcome of mutations in *ARX gene* were predicted using several computational analysis tools. Up to our knowledge, this is the first in silico study in the coding region of *ARX* gene which aims to classify the most damaging nsSNPs in this gene, that might be useful for further analysis and testing in therapeutic drug clinical trials.

## 2 Materials and Methods

### 2.1 Data mining

The raw data of *ARX* gene were recovered from NCBI website [16]; while the protein reference sequence was retrieved from Uniprot [17].

### 2.2 Functional Analysis

#### 2.2.1 SIFT

SIFT was used to discern the effect of each amino acid substitution on *ARX* protein function. SIFT predicts damaging SNPs on the basis of the degree of conserved amino acid residues in aligned sequences to the closely related sequences, assembled through PSI-BLAST [18].

#### 2.2.2. PolyPhen-2

PolyPhen 2 sever (Polymorphism Phenotyping v2), was used to study the potential effects of each amino acid substitution on the structural and functional properties of *ARX* protein by applying physical and comparative approaches [19].

#### 2.2.3. PROVEAN

PROVEAN is a trained functional online tool that depends on alignment-based score. Mutations are predicted to have neutral or deleterious effects based on their alignment score being more or less than -2.5 respectively. [20].

#### 2.2.4. SNAP2

SNAP2 is a trained functional tool that is based on neural network. It differentiates between disease and benign mutations by taking a variety of features into account. it got an accuracy of 83% [21].

#### 2.2.5. SNPs&GO

It is a support vector machine (SVM) based on the method to accurately predict the disease-related mutations from protein sequence. The probability score higher than 0.5 reveals the disease-related effect of mutation on the parent protein function. [22]. SNPs&GO has another tool impeded with it called PHD-SNP which classifies the results into deleterious or neutral mutation [23].

#### 2.2.6. P-Mut

P-Mut is a functional web-based tool which permits swift and accurate results (80%) of the compulsive features of each SNP based on neural networks intelligence [24].

#### 2.2.8. Condel

Condel is a method to assess the outcome of non-synonymous single nucleotide variants SNVs using consensus deleteriousness score that combines various tools (SIFT, Polyphen2, Mutation Assessor, and FATHMM). The result of the prediction are either deleterious or neutral with a score ranging from zero to one [25].

### 2.3. Stability analysis

#### 2.3.1. I-Mutant 3.0

I-Mutant 3.0 is a SVM based tool, it predicts whether a SNPs mutation stabilizes or destabilizes the protein structure by calculating free energy change through coupling predictions with the energy based FOLD-X tool [26].

#### 2.3.2. MUPro

MUPro is a structural analysis online tool for the calculation of protein stability variations upon arbitrarily SNPs. A score < 0 means that the mutant decreases the protein stability; conversely, a score > 0 means the mutant increases the protein stability [27].

### 2.4. ClinVar

ClinVar is an archive for evidence-based studies of the associations among human mutations and phenotypes. It was used to associate our predicted results with the possible clinical one [28]

### 2.5. Variant Effect Predictor (VEP)

Variant Effect Predictor provides toolset to annotate and assist in prioritization of variants in both large-scale sequencing projects and smaller analysis studies. The SNPs IDs for the SNPs of interest was retrieved from dbSNP database, which was then used as an input to predict the Functional consequences of these variants [29].

### 2.6. RaptorX

RaptorX have been used to generate a 3D structural model for wildtype ARX. The FASTA format sequence of *ARX* protein was retrieved from UniProt, it was then used as an input to predict the 3D structure of human *ARX* protein [30].

### 2.7. UCSF Chimera

UCSF Chimera is a visualization tool of 3D structure prototype. A predicted model was formed by RaptorX to visualize and compare the wild and mutant amino acids using UCSF Chimera [31].

### 2.8. Project HOPE

Project HOPE is a server to search protein 3D structures by data mining from several sources databases. it was used to analyze the physiochemical changes induced by the mutations and to confirm the outcomes that we achieved earlier [32].

### 2.9. Mutation3D

Mutation3D is a trained web server, offers driver genes in cancer by identifying clusters of amino acid substitutions within tertiary protein structures. The web interface allows analyzing mutations in a variety of formats and provides unified access to explore mutation clusters derived from over 975,000 somatic mutations reported by 6,811 cancer sequencing studies [33].

### 2.10. SOPMA

SOPMA can predict the secondary structure of 69.5% of amino acids in the entire database containing 126 non-homologous protein chains. When SOPMA and a neural network method (PHD) are joint correctly they can predicts 82.2% of residues for 74% of co-predicted amino acids [34].

### 2.11. Search Tool for the Retrieval of Interacting Genes (STRING)

Search Tool for the Retrieval of Interacting Genes (STRING) was applied to identify the interactions of *ARX* protein with other corresponding proteins [35]. This server was used to predict the interaction of human *ARX* protein with other proteins by providing accurate prediction and experimental interaction data of *ARX*.

### 2.12. PSIPRED

Protein secondary structures was predicted based on their specific matrices using PSIPRED server, with matrices developed by PSI-BLAST [36].

### 2.13. GeneMANIA

GeneMANIA is an approach to recognize protein function by prediction integrating multiple genomics and proteomics sources to make inferences about the function of unknown proteins [37].

## 3. Results

The total number of SNPs in the coding region that were recovered from NCBI was 273 nsSNPs. These SNPs were submitted into different functional analysis online tools **(Figure 1)**. 213 out of 273 nsSNPs were found to be affected by SIFT, 143 damaging SNPs (5 possibly damaging and 138 probably damaging) by Pholyphen-2, 193 were found to be deleterious by PROVEAN and 179 were predicted damaging by SNAP2 **(Table 1)**. After another filtration, the number of SNPs decreased to 19 and then were submitted into SNPs&GO, PHD-SNP, P-Mut and Condel to give more accurate results on their effects on the functional impact; the quad positive from the four tools were 11 SNPs **(Table 2; Figure 3)**. The stability analysis on these 12 SNPs was tested by I-Mutant3.0 and MUPro; which discovered that all 11 SNPs decrease the protein stability **(Table 3)**. The secondary structure of ARX was predicted by SOPMA, which explained the distributions of alpha helix, beta sheet, and coil. The result indicated a large number of random coils (257, 45.73%), followed by 245 alpha helixes (43.59%), 33 extended strands (5.87%) and 27 beta turns (4.80%) in the predicted secondary structure **(Figure 6, 7)**.

**Table 1:** Deleterious SNPs in *ARX* gene as predicted by four in silico tools:

| SUB | SIFT prediction | Score | Polyphen2 prediction | Score | PROVEAN prediction | Score | SNAP2 prediction | Score |
| --- | --- | --- | --- | --- | --- | --- | --- | --- |
| L535Q | Deleterious | 0 | probably damaging | 1 | Deleterious | -5.204 | Deleterious | 54 |
| R528S | Deleterious | 0 | probably damaging | 1 | Deleterious | -4.727 | Deleterious | 57 |
| G461D | Deleterious | 0 | probably damaging | 1 | Deleterious | -4.066 | Deleterious | 52 |
| P420L | Deleterious | 0 | probably damaging | 1 | Deleterious | -5.372 | Deleterious | 1 |
| R384H | Deleterious | 0 | probably damaging | 1 | Deleterious | -4.481 | Deleterious | 57 |
| R380L | Deleterious | 0 | probably damaging | 1 | Deleterious | -6.341 | Deleterious | 82 |
| V374D | Deleterious | 0 | probably damaging | 1 | Deleterious | -6.507 | Deleterious | 79 |
| L343Q | Deleterious | 0 | probably damaging | 1 | Deleterious | -4.641 | Deleterious | 63 |
| T333N | Deleterious | 0 | probably damaging | 1 | Deleterious | -3.734 | Deleterious | 80 |
| T333S | Deleterious | 0 | probably damaging | 1 | Deleterious | -3.17 | Deleterious | 67 |
| R332H | Deleterious | 0 | probably damaging | 1 | Deleterious | -3.767 | Deleterious | 79 |
| R330H | Deleterious | 0 | probably damaging | 1 | Deleterious | -3.8 | Deleterious | 52 |
| R57H | Deleterious | 0 | probably damaging | 1 | Deleterious | -2.595 | Deleterious | 71 |
| R42L | Deleterious | 0 | probably damaging | 1 | Deleterious | -3.633 | Deleterious | 89 |
| R42W | Deleterious | 0 | probably damaging | 1 | Deleterious | -4.152 | Deleterious | 93 |
| P38L | Deleterious | 0 | probably damaging | 1 | Deleterious | -5.524 | Deleterious | 28 |
| G34R | Deleterious | 0 | probably damaging | 1 | Deleterious | -4.419 | Deleterious | 41 |
| L33P | Deleterious | 0 | probably damaging | 1 | Deleterious | -3.867 | Deleterious | 73 |
| D30Y | Deleterious | 0 | probably damaging | 1 | Deleterious | -4.738 | Deleterious | 86 |

**Table 2:** The most damaging SNPs in ARX gene as predicted by four different online tools:

| SUB | SNPs&GO Prediction | RI | Probability | PHD-SNP Prediction | RI | Probability | P-mut Prediction | Probability | CONDEL Prediction | Score |
| --- | --- | --- | --- | --- | --- | --- | --- | --- | --- | --- |
| L535Q | Deleterious | 4 | 0.706 | Deleterious | 6 | 0.776 | Disease | 0.73 | Deleterious | 0.77 |
| R528S | Deleterious | 1 | 0.549 | Deleterious | 2 | 0.577 | Disease | 0.7 | Deleterious | 0.58 |
| G461D | Deleterious | 5 | 0.727 | Deleterious | 4 | 0.704 | Neutral | 0.43 | Deleterious | 0.76 |
| R380L | Deleterious | 5 | 0.738 | Deleterious | 7 | 0.84 | Disease | 0.82 | Deleterious | 0.70 |
| V374D | Deleterious | 6 | 0.792 | Deleterious | 8 | 0.891 | Disease | 0.83 | Deleterious | 0.78 |
| L343Q | Deleterious | 6 | 0.78 | Deleterious | 7 | 0.863 | Disease | 0.83 | Deleterious | 0.69 |
| T333N | Deleterious | 3 | 0.658 | Deleterious | 6 | 0.806 | Disease | 0.79 | Deleterious | 0.64 |
| T333S | Deleterious | 2 | 0.613 | Deleterious | 3 | 0.655 | Disease | 0.75 | Deleterious | 0.71 |
| R332H | Deleterious | 6 | 0.785 | Deleterious | 7 | 0.83 | Disease | 0.76 | Deleterious | 0.68 |
| R330H | Deleterious | 6 | 0.778 | Deleterious | 6 | 0.808 | Disease | 0.73 | Deleterious | 0.54 |
| G34R | Deleterious | 1 | 0.543 | Deleterious | 3 | 0.653 | Disease | 0.53 | Deleterious | 0.53 |
| L33P | Deleterious | 3 | 0.635 | Deleterious | 4 | 0.722 | Disease | 0.53 | Deleterious | 0.53 |

**Table 3:**
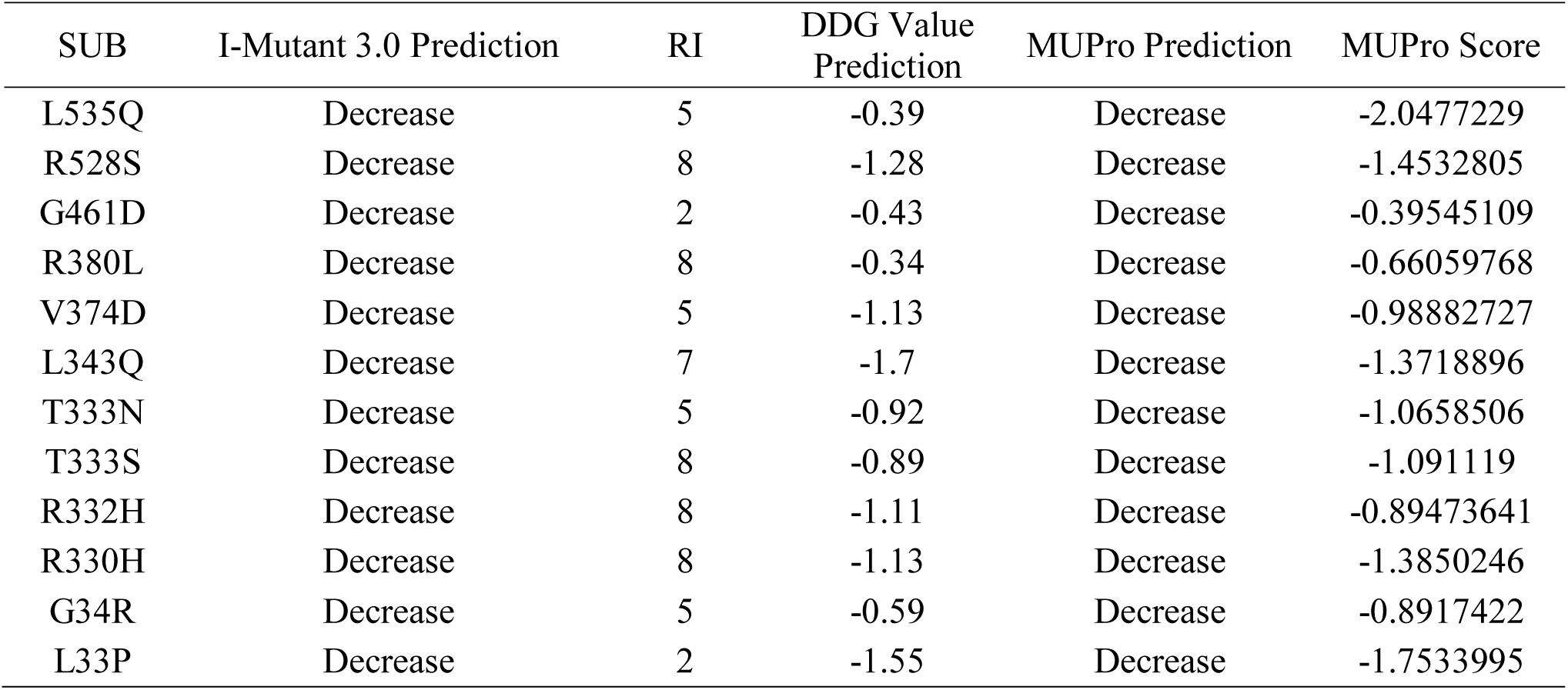
Protein stability prediction on ARX protein, investigated using I-Mutant 3.0 and MUPro:

**Table 4:** Shows variant consequences, transcripts, and regulatory features by VEP tool:

| rs variants | Consequence | Impact | Feature type | Biotype |
| --- | --- | --- | --- | --- |
| rs387906715 | Missense variant | Moderate | Transcript | Protein coding |
| rs387906715 | Downstream gene variant | Modifier | Transcript | Processed transcript |
| rs387906715 | Downstream gene variant | Modifier | Transcript | Protein coding |
| rs387906715 | Regulatory region variant | Modifier | Regulatory Feature | Protein coding |
| rs1457617771 | Missense variant | Moderate | Transcript | Protein coding |
| rs1457617771 | Downstream gene variant | Modifier | Transcript | Processed transcript |
| rs1457617771 | Downstream gene variant | Modifier | Transcript | Protein coding |
| rs1457617771 | Regulatory region variant | Modifier | Regulatory Feature | Protein coding |
| rs1195367604 | Missense variant | Moderate | Transcript | Protein coding |
| rs1195367604 | Upstream gene variant | Modifier | Transcript | Processed transcript |
| rs1195367604 | Upstream gene variant | Modifier | Transcript | Protein coding |
| rs1195367604 | Regulatory region variant | Modifier | Regulatory Feature | Protein coding |
| rs867820539 | Missense variant | Moderate | Transcript | Protein coding |
| rs867820539 | Upstream gene variant | Modifier | Transcript | Processed transcript |
| rs867820539 | Upstream gene variant | Modifier | Transcript | Protein coding |
| rs867820539 | Regulatory region variant | Modifier | Regulatory Feature | Protein coding |
| rs587783183 | Missense variant | Moderate | Transcript | Protein coding |
| rs587783183 | Upstream gene variant | Modifier | Transcript | Processed transcript |
| rs587783183 | Upstream gene variant | Modifier | Transcript | Protein coding |
| rs587783183 | Regulatory region variant | Modifier | Regulatory Feature | Protein coding |
| rs104894741 | Missense variant | Moderate | Transcript | Protein coding |
| rs104894741 | Downstream gene variant | Modifier | Transcript | Processed transcript |
| rs104894741 | Downstream gene variant | Modifier | Transcript | Processed transcript |
| rs104894745 | Missense variant | Moderate | Transcript | Protein coding |
| rs104894745 | Missense variant | Moderate | Transcript | Protein coding |
| rs104894745 | Downstream gene variant | Modifier | Transcript | Processed transcript |
| rs104894745 | Downstream gene variant | Modifier | Transcript | Processed transcript |
| rs104894745 | Downstream gene variant | Modifier | Transcript | Processed transcript |
| rs104894745 | Downstream gene variant | Modifier | Transcript | Processed transcript |
| rs111033612 | Missense variant | Moderate | Transcript | Protein coding |
| rs111033612 | Missense variant | Moderate | Transcript | Protein coding |
| rs111033612 | Downstream gene variant | Modifier | Transcript | Processed transcript |
| rs111033612 | Downstream gene variant | Modifier | Transcript | Processed transcript |
| rs111033612 | Downstream gene variant | Modifier | Transcript | Processed transcript |
| rs111033612 | Downstream gene variant | Modifier | Transcript | Processed transcript |
| rs886039308 | Missense variant | Moderate | Transcript | Protein coding |
| rs886039308 | Downstream gene variant | Modifier | Transcript | Processed transcript |
| rs886039308 | Downstream gene variant | Modifier | Transcript | Processed transcript |
| rs1445093587 | Missense variant | Moderate | Transcript | Protein coding |
| rs1445093587 | Non-coding transcript exon variant | Modifier | Transcript | Processed transcript |
| rs1445093587 | Non-coding transcript exon variant | Modifier | Transcript | Processed transcript |
| rs1445093587 | Regulatory region variant | Modifier | Regulatory Feature | Processed transcript |
| rs28936077 | Missense variant | Moderate | Transcript | Protein coding |
| rs28936077 | Non-coding transcript exon variant | Modifier | Transcript | Processed transcript |
| rs28936077 | Non-coding transcript exon variant | Modifier | Transcript | Processed transcript |
| rs28936077 | Regulatory region variant | Modifier | Regulatory Feature | Processed transcript |

**Figure 1:**
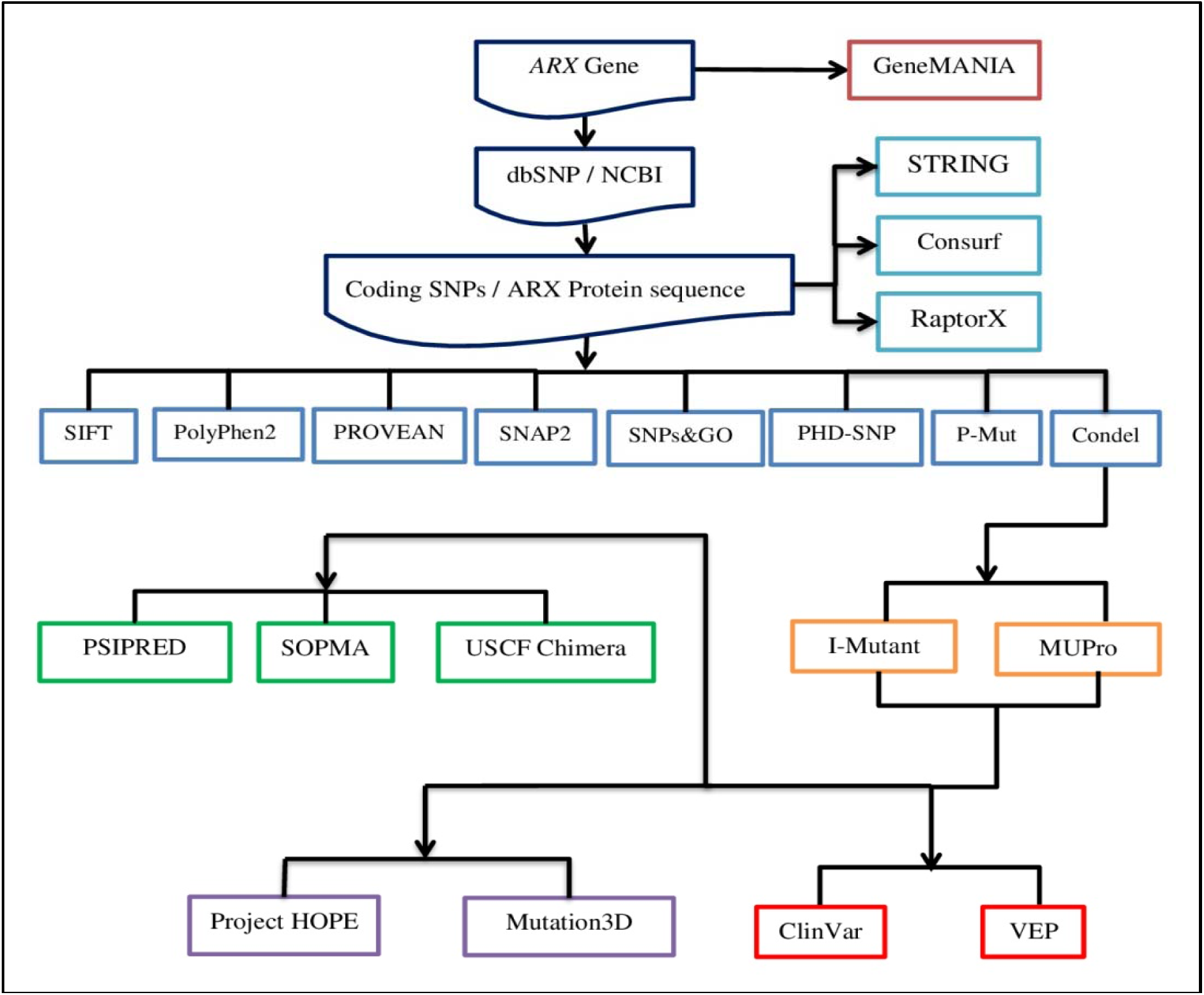
Schematic representation of *in silico* tools for computational analysis of *ARX*.

**Figure 2:**
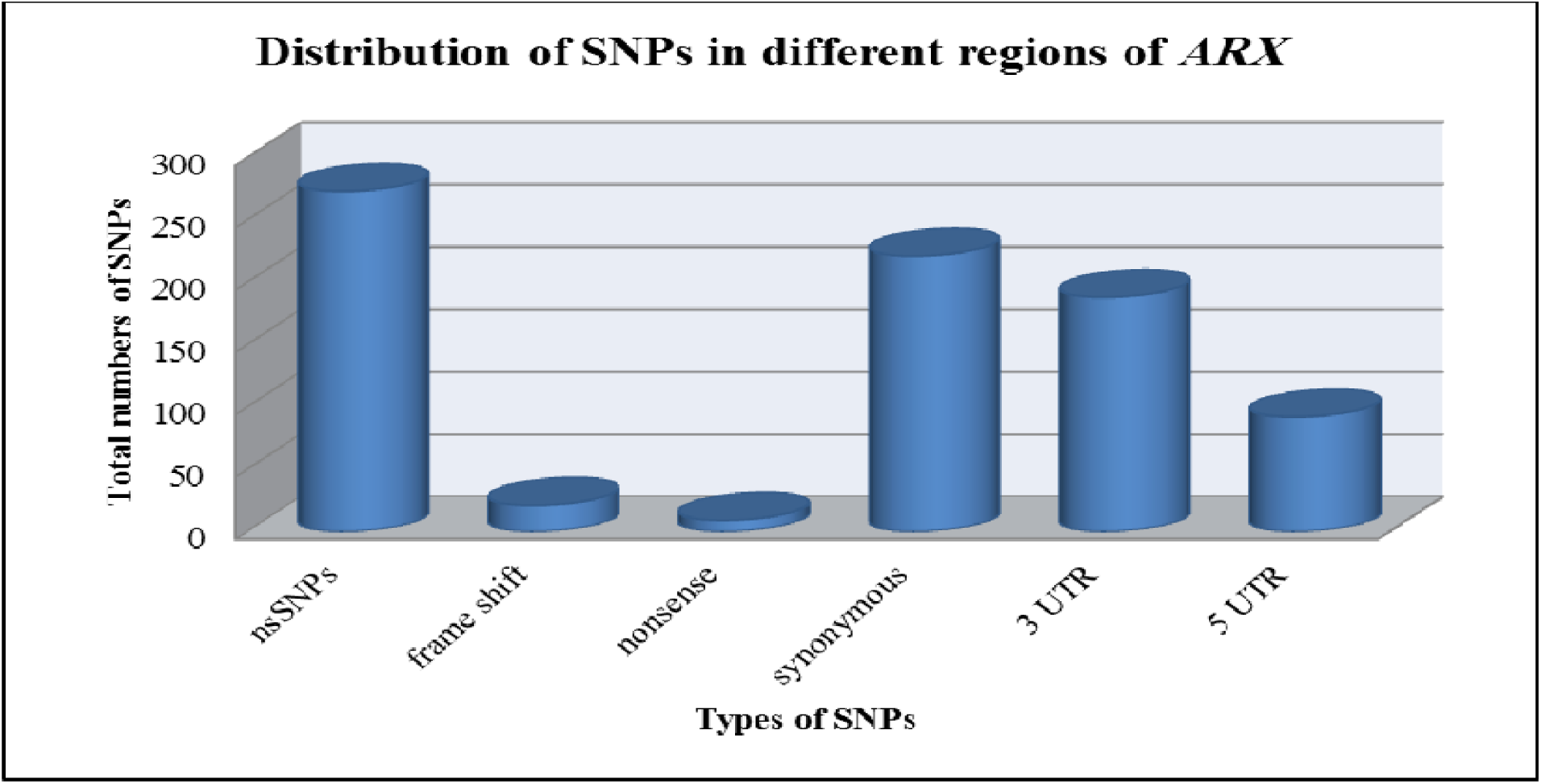
Distribution of SNPs in different regions of *ARX* gene.

**Figure 3:**
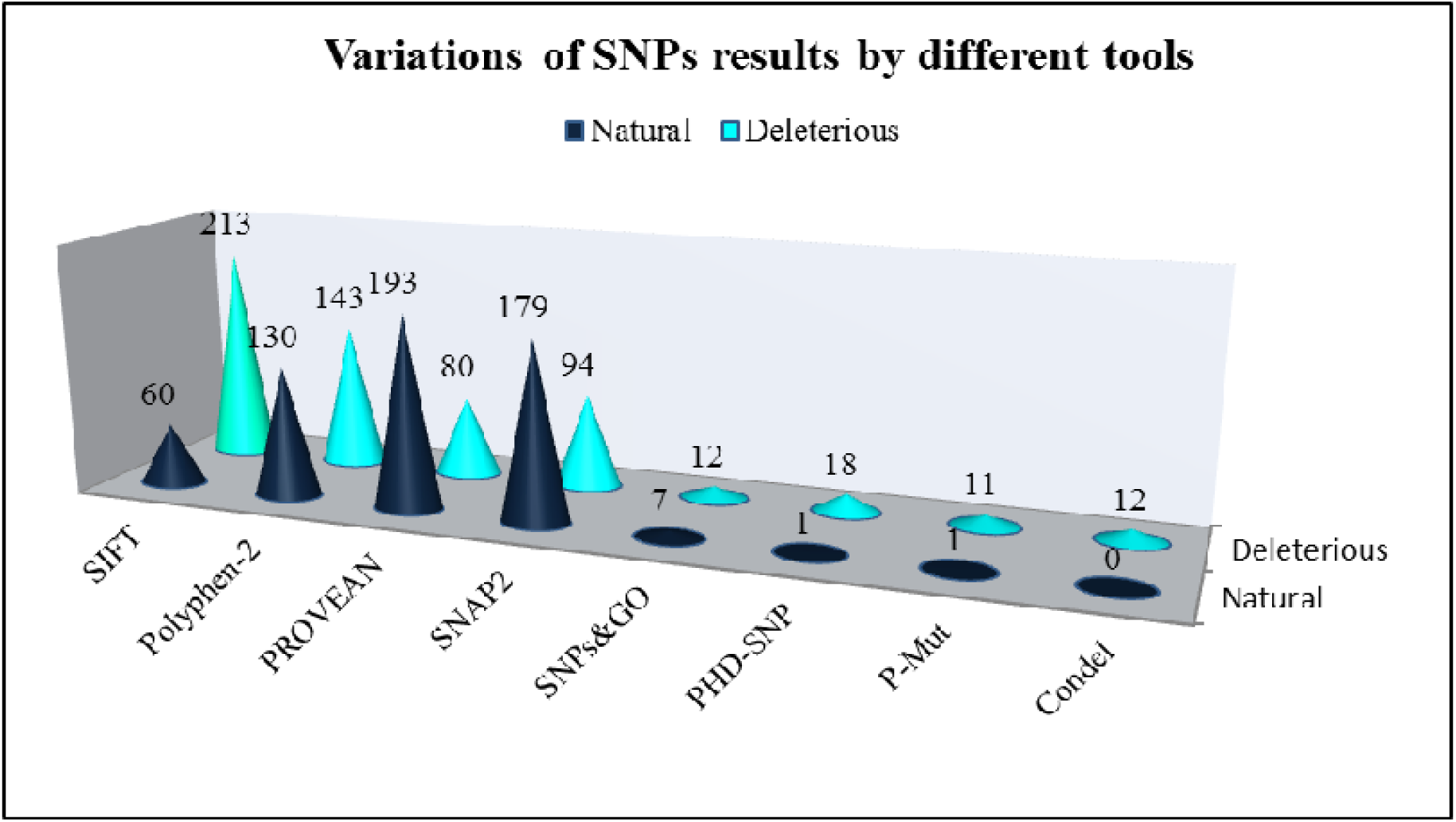
Variations of SNPs results in ARX gene as predicted by eight different tools.

**Figure 4:**
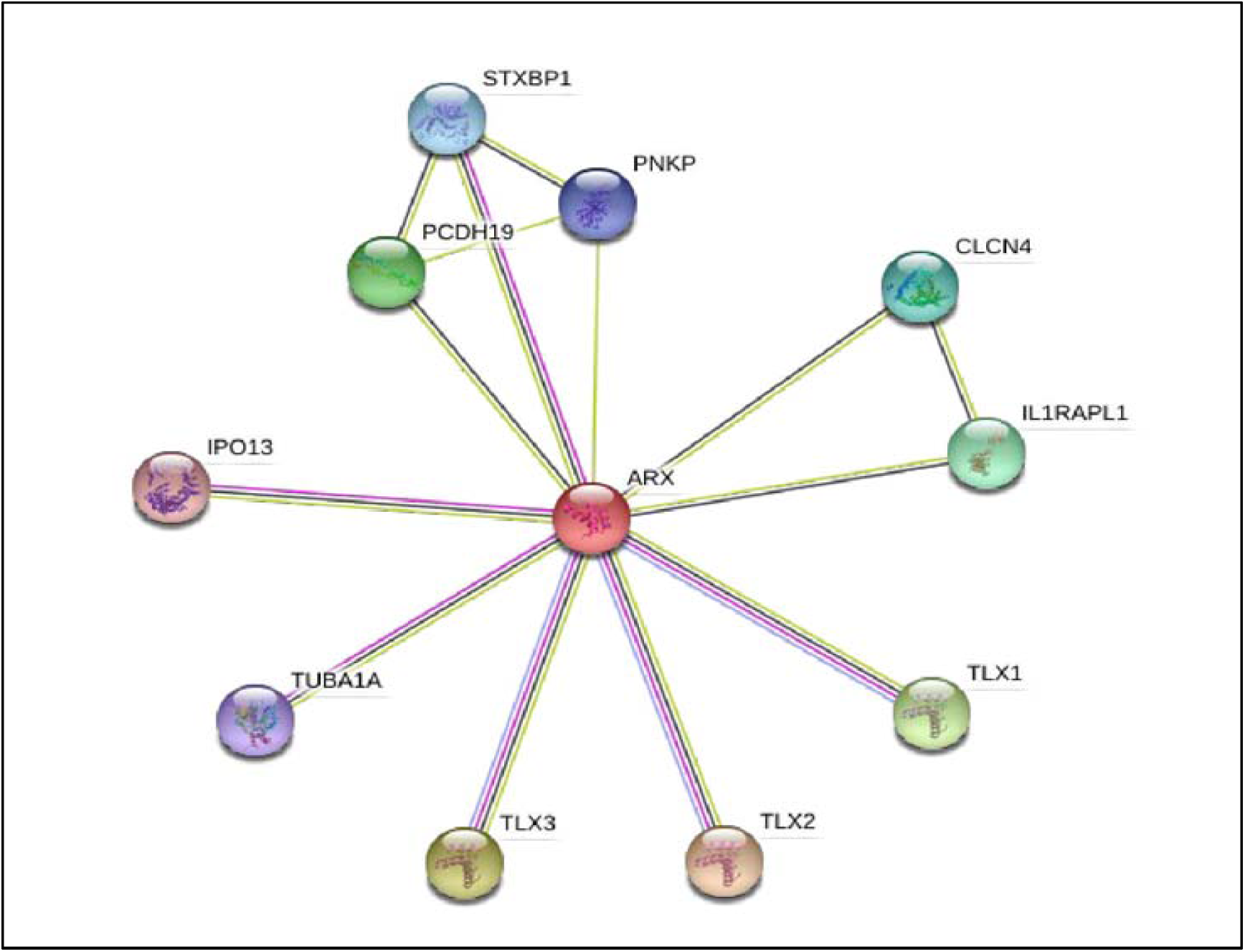
Protein–protein interaction network of ARX protein shown by STRING.

**Figure 5:**
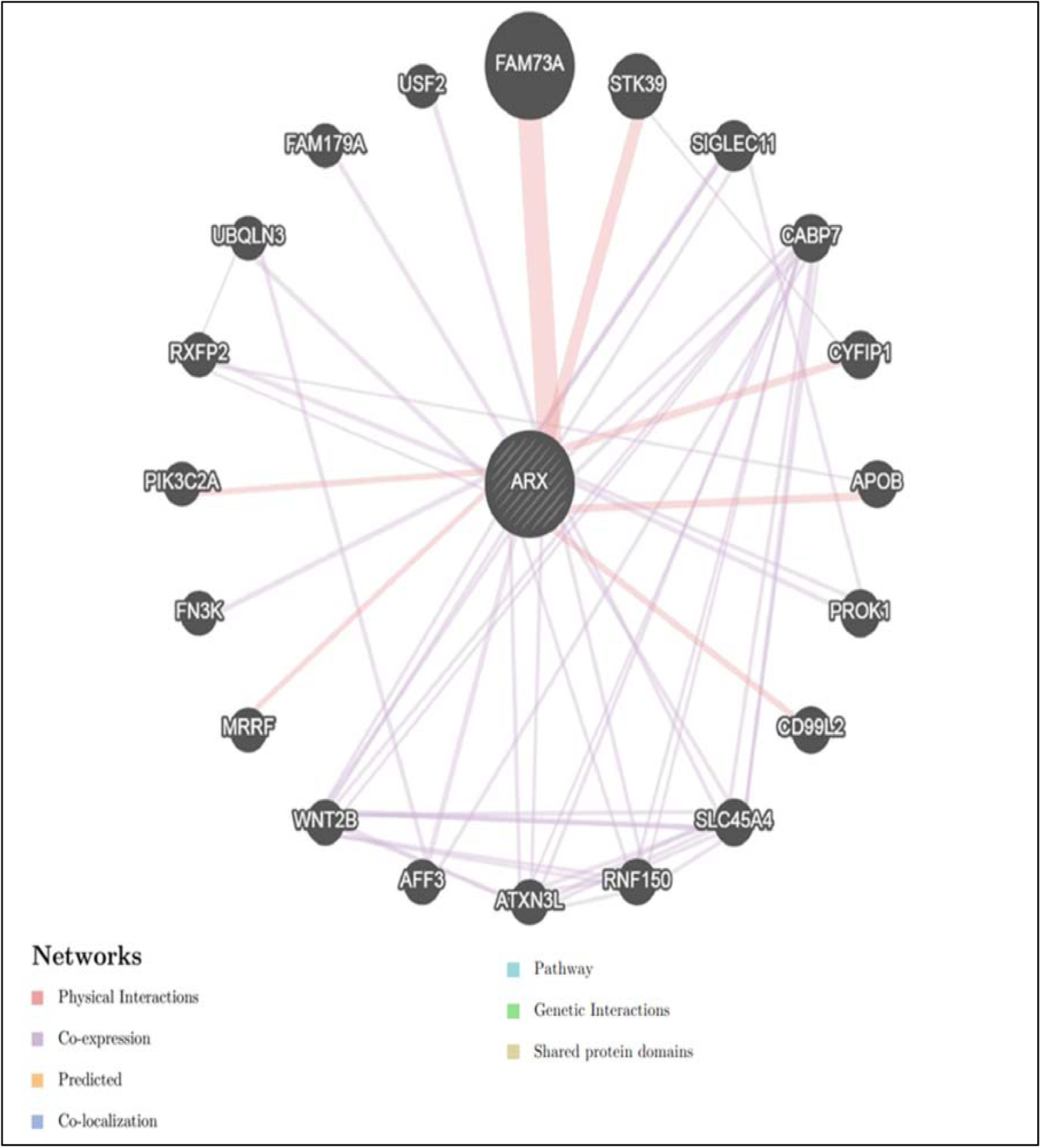
Biological network Interaction between *ARX* and its related genes, as predicted by GeneMANIA

**Figure 6:**
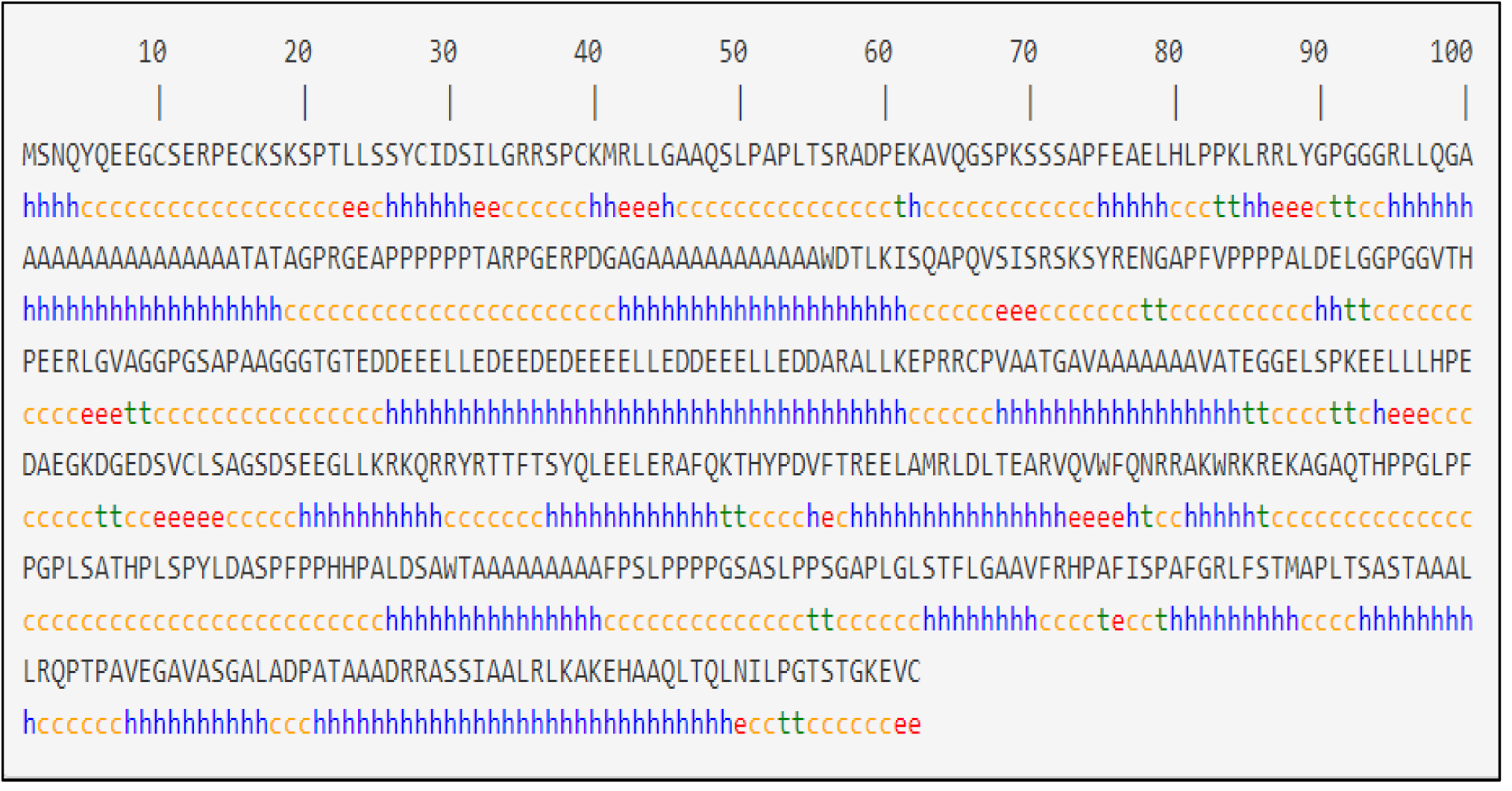
SOPMA analysis of the secondary structure of individual amino acid residues in protein produced from the ARX gene.

**Figure 7:**
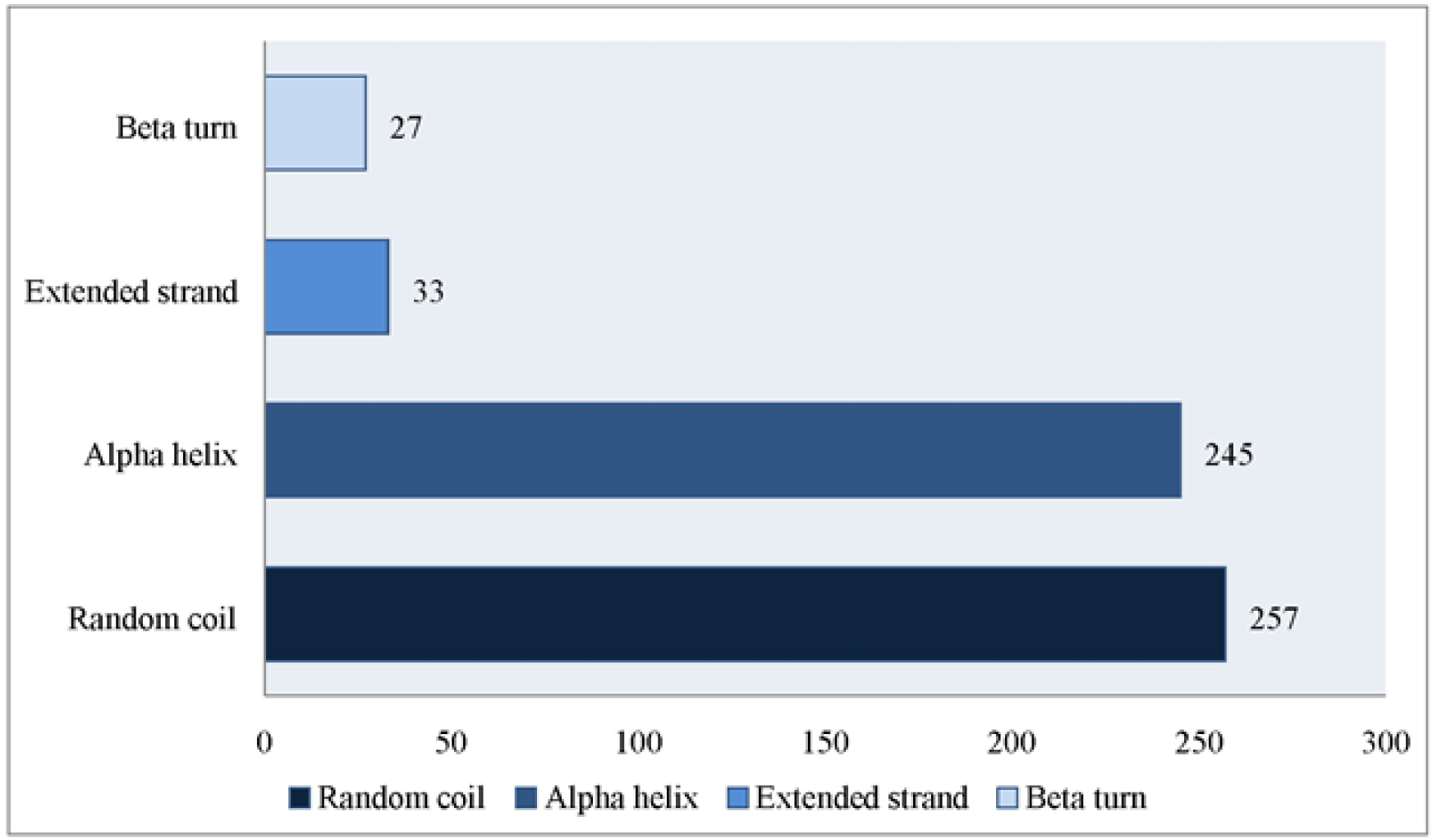
Distribution of high-risk nsSNPs of ARX in random coils, alpha helixes, extended strands and beta turns predicted by SOPMA.

## 4. Discussion

Disease-causing SNPs are frequently found to arise at evolutionarily conserved regions. Those have a critical role at structural and functional levels of the protein [38-40]. Therefore, our effort was devoted to the coding region that is evolutionary conserved in most cases, which revealed 11 mutations in *ARX* gene as predicted by different in *silico* tools **(Figure 1)**.

The 11 pathogenic SNPs come after extensive *in silico* analysis; eight tools **(Figure 2)** were used to investigate the effects of each SNP on the functional impact; the reason why results are different in each software are due to the different associated algorithm of these programs **(Tables 1 and 2)**. A silent mutation may alter sequences involved in splicing and lead to abnormal transcripts. Such mutations risk being overlooked; an example to take into account is that of an apparently silent mutation on the *ARX* gene (G461D in **Table 2**). On the other hand, Structural Consequences of these 11 SNPs were studied by I-Mutant3.0 and MUPro tools, the analysis revealed that all 12 SNPs decrease the protein stability by both tools **(Table 3)** thus proposing that these variants could destabilize the amino acid interactions causing functional deviations of protein to some point.

All these SNPs (R528S, R380L, T333S, and G34R) were retrieved from the dbSNP as untested and all were found to be deleterious mutations; while these SNPs (L535Q, T333N, R332H and L33P) were recovered as deleterious which agrees with our outcomes **(Table 2)**.

STRING database predicted the functional interaction pattern of ARX protein to other proteins in a cell. Strong functional associations of ADIPOR1 protein have been observed with IPO13, TUBA1A, TLX3, TLX2, TLX1 and STXBP1 partners. Besides, weak interactions with less confidence have been observed for PNKP protein; this draws attention to the consequences of ARX mutations could affect other proteins involved in central nervous systems function **(Figure 4)**.

We also used GeneMANIA sever, which showed that *ARX* has many vital functions, which if disturbed may leads to neuroblastoma [10]. Furthermore, any disturbance in ARX function is thought to disrupt the normal brain development, leading to seizures and intellectual disability [11, 12]. The genes co-expressed with, sharing similar protein domain, or contributed to achieve similar function as shown in **(Figure 5)**.

According to SOPMA secondary structure calculations, 11 high-risk nsSNPs were located in 10 amino acid sites, and 89.32% of sites were located in alpha helixes and random coils, which is in accordance with the evidence from literature that both deleterious and polymorphic mutations are mainly located in helixes and coil regions and not frequently in β turns [41] **(Figure 6,7)**.

We also used PSIPRED to predict Secondary structure of *ARX* protein ; the results highlight a mix distribution of coil and alpha helix. The coil was observed to be the major secondary structural motif (45.73%), followed by helix (43.59%) as generated by PSIPRED. However, it is worthy to note that protein-protein interaction occurs at the level of protein tertiary structure and this makes it important to predict the impact of nsSNPs on tertiary structure of *ARX* gene. Building on these predictions we can assume that these pathogenic mutations in the human *ARX gene* could result in altered stability and metal binding, as well as loss of allosteric and catalytic residue with a concomitant loss of enzyme activity. Furthermore, we used the STRING maps to illustrate the protein–protein interaction of *ARX* **(Figure 8)**. This is useful in understanding the genotype-phenotype consequence of the mutation [42].

**Figure 8:**
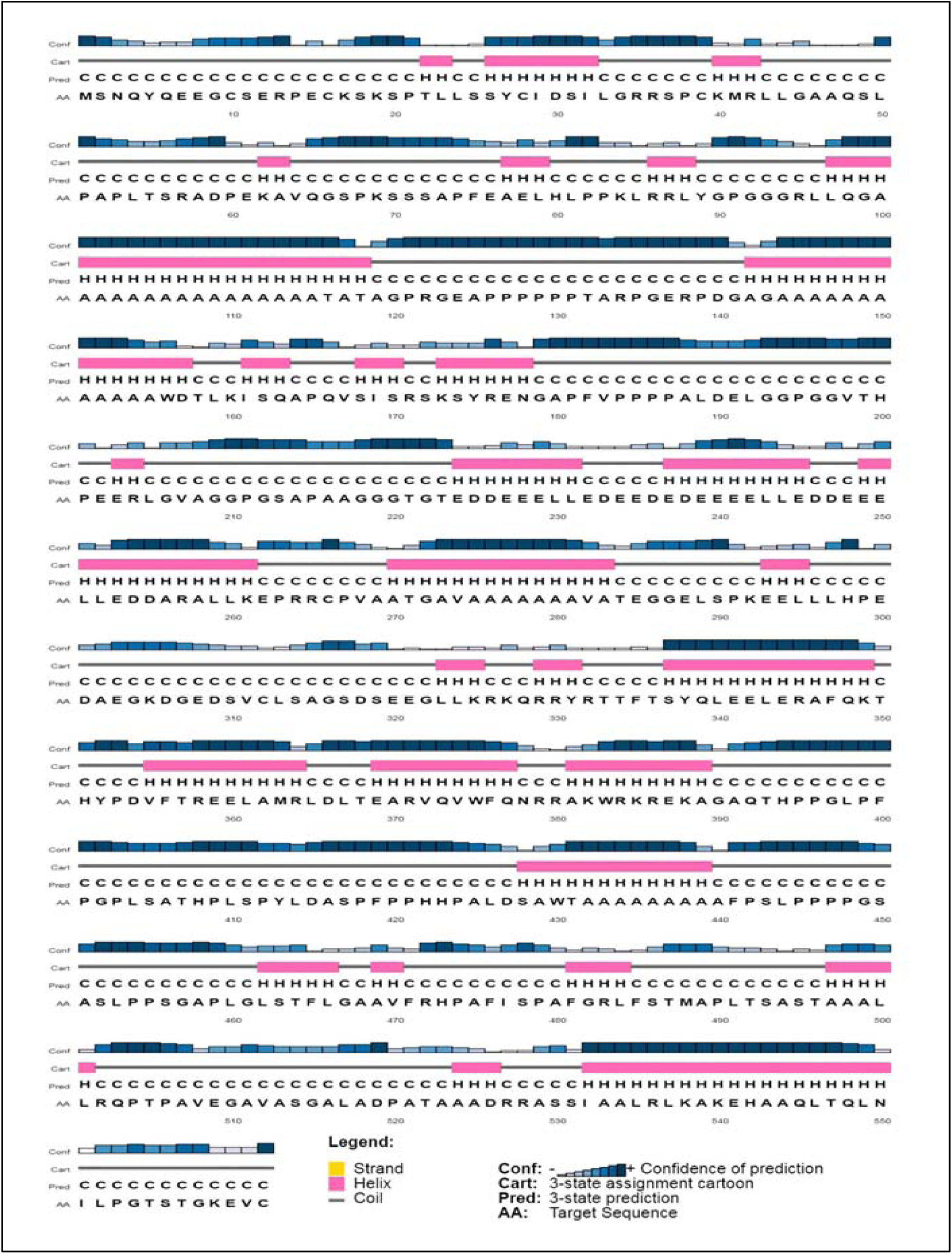
Secondary structure of ARX showing beta helix and coil. The results highlight a mix distribution of coil and alpha helix, as generated by PSIPRED.

We also used Consurf server to check the conservation region of ARX protein, and the result shows 9 SNPs (L33P, G34R, R332H, T333S, T333N, V374D, R380L, R528S and L535Q) located in highly conserved regions, which can directly affect the protein function **(Figure 9)**.

**Figure 9:**
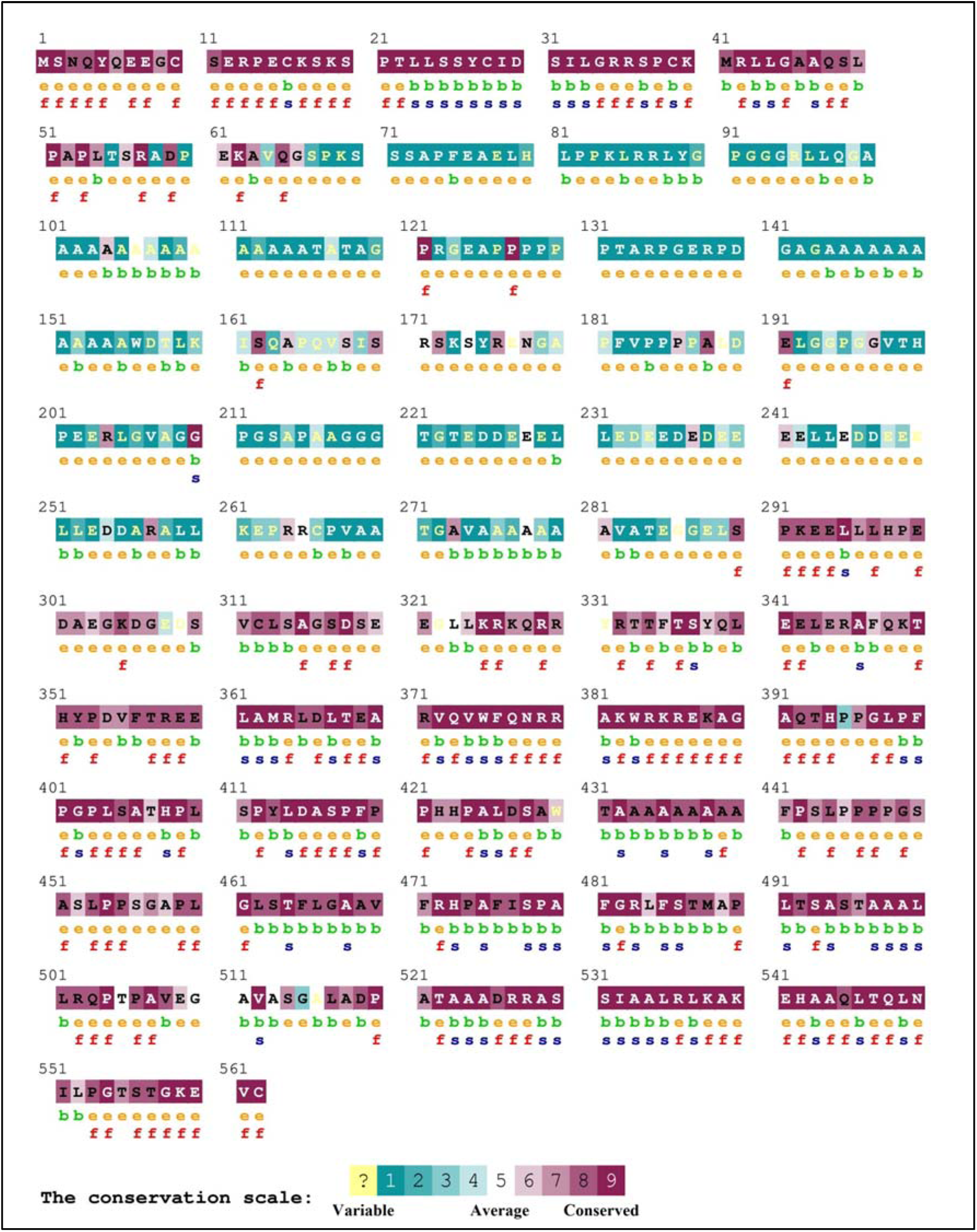
The conserved amino acids across species in ARX protein were determined using ConSurf.

In order to study the biophysical properties of these mutations, Project HOPE server was used to serve this purpose; RaptorX was used to predict a 3D structure model for ARX protein, while UCSF Chimera was used to visualize the amino acids change **(Figure 10)**. In **(Figure 11)**: (R380L): the amino acid Arginine changes to Leucine at position 380; the residue is located in a DNA binding region. The differences in properties between wild-type and mutant residue can easily cause loss of these interactions or disturb the domain which will affect the function of the protein. The wild-type residue forms a hydrogen bond with Histidine at position 351. T\due to the size difference between the wild-type and mutant residue the mutation will cause an empty space in the core of the protein affecting its function.

**Figure 10:**
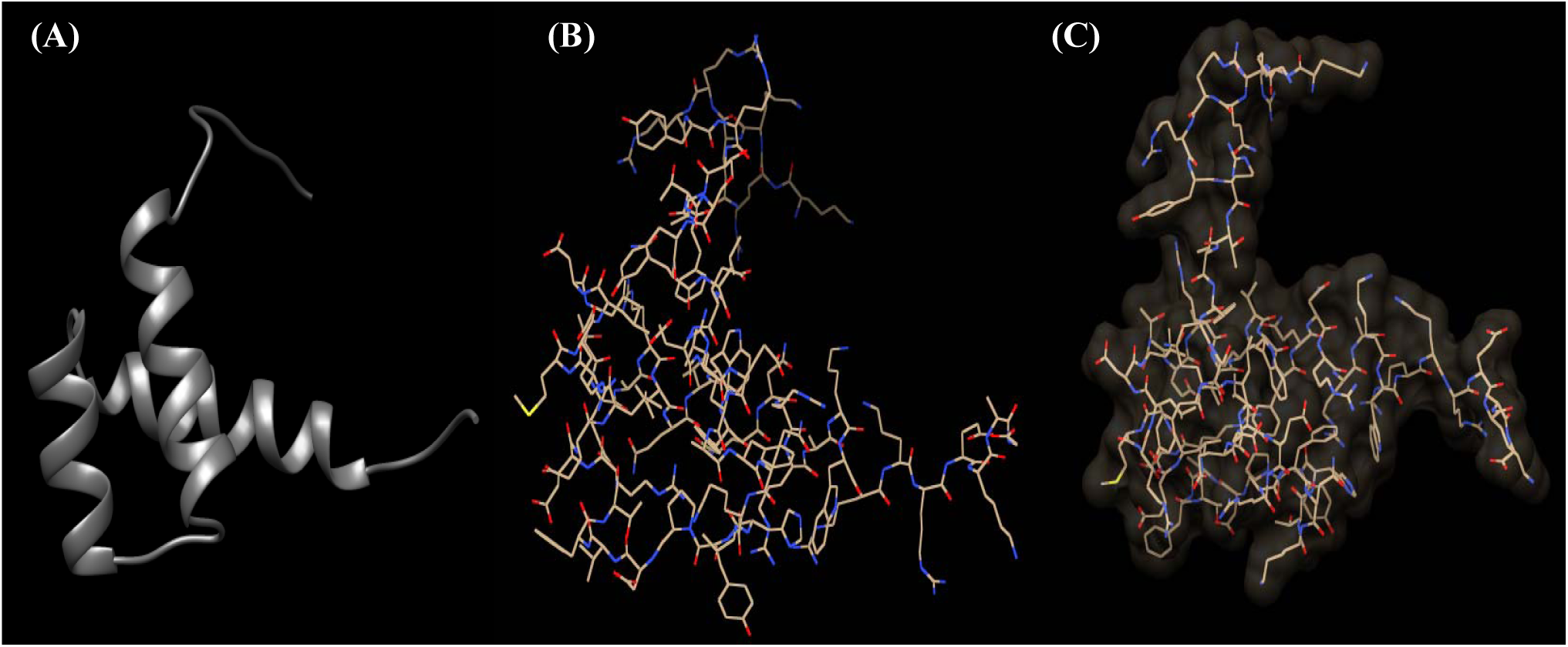
The 3D structure of ARX protein models by (A) RaptorX, (B) CPHmodels-3.2 and (C) SWISS-MODEL Servers. The 3D structure of the ARX protein model was created by using RaptorX; it could not create the 3D model of all amino acid positions. Therefore, the model was done from position 325 to 390, due to the lack of data.

**Figure 11:**
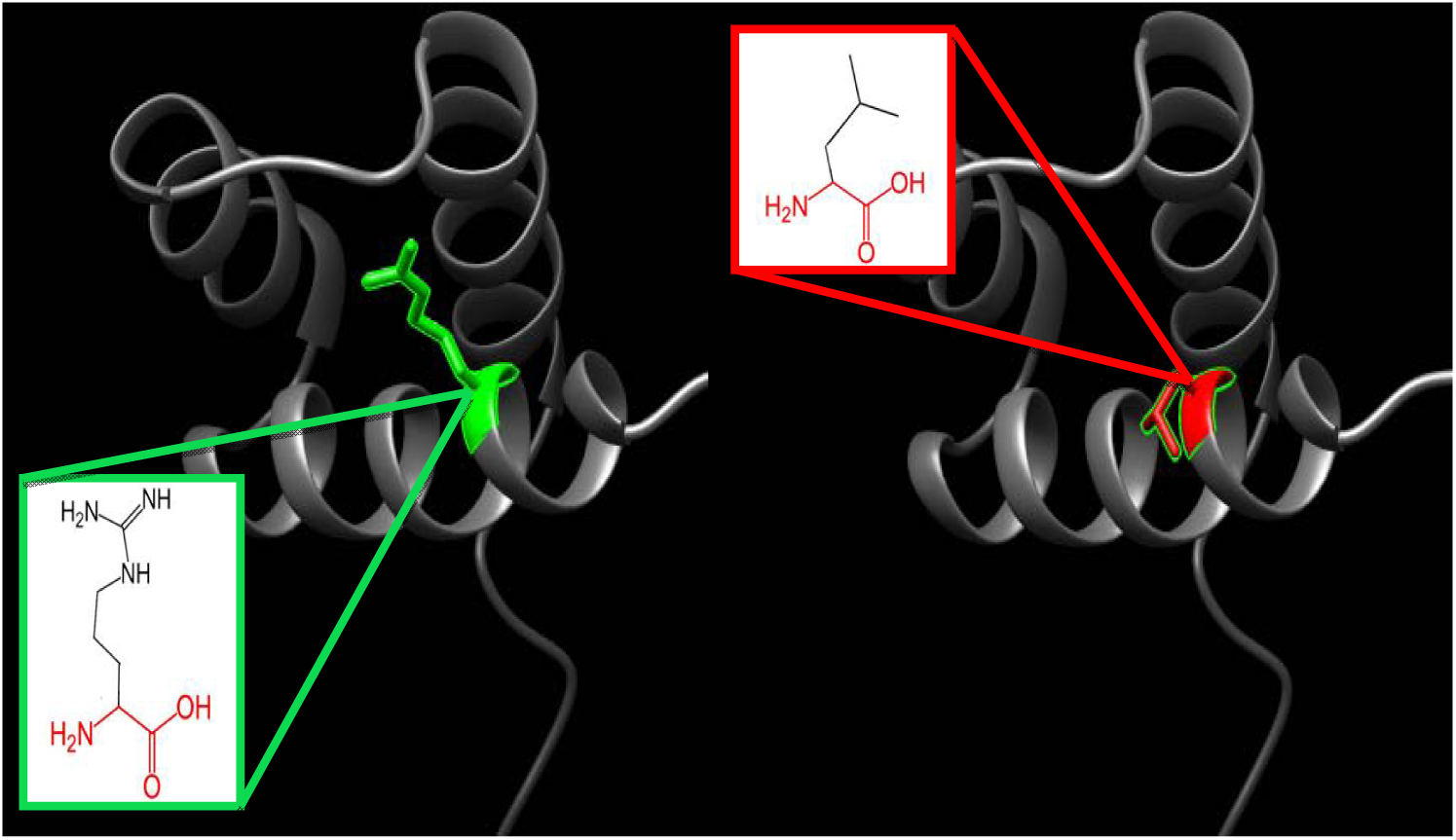
(R380L): the amino acid Arginine changes to Leucine at position 380. This visualization was done using Chimera software coupled with the 2D structures (in boxes) from Project Hope report of ARX gene mutations. The green color indicates the wild amino acid, and the red color indicates the mutant one, while the gray color indicates the rest of ARX protein structure.

In **(Figure 12)**: (V374D): the amino acid Valine changes to Aspartic Acid at position 374; the residue has DNA interactions. The mutation introduces a charge at this position; this can cause repulsion between the mutant residue and neighboring residues. The residue is located on the surface of the protein; hence mutation of this residue can disturb interactions with other molecules or other parts of the protein. The difference in hydrophobicity of the Valine and Aspartic acid residue might lead to loss of hydrophobic interactions with other molecules on the surface of the protein.

**Figure 12:**
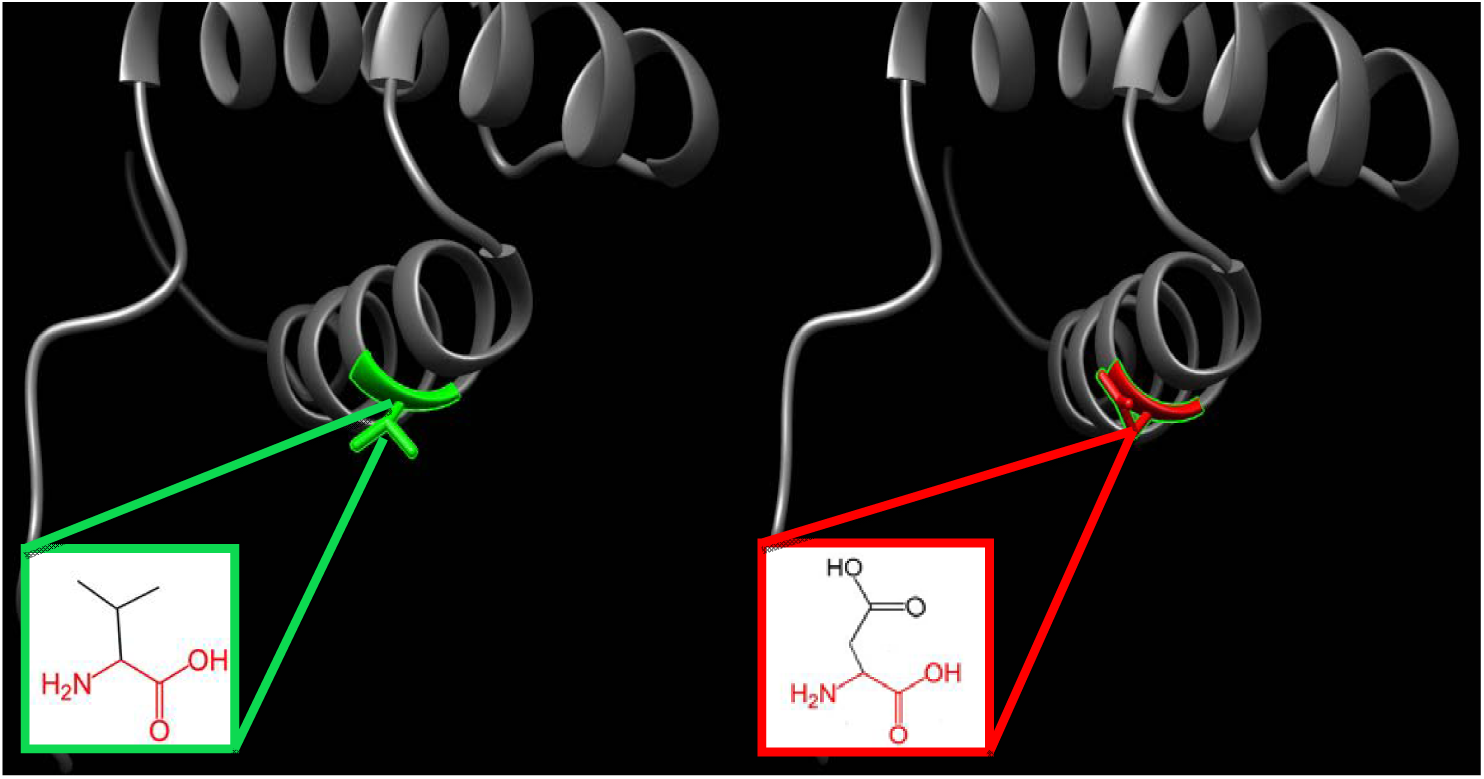
(V374D): the amino acid Valine changes to Aspartic Acid at position 374. This visualization was done using Chimera software coupled with the 2D structures (in boxes) from Project Hope report of ARX gene mutations. The green color indicates the wild amino acid, and the red color indicates the mutant one, while the gray color indicates the rest of ARX protein structure.

In **(Figure 13)**: (L343Q): the amino acid Leucine changes to Glutamine at position 343; in the 3D-structure, it can be seen that the wild-type residue is located in an α-helix. The mutation converts the wild-type residue to Glutamine which does not prefer α-helices as secondary structure. The residue is found on the surface of the protein, mutation of this residue can disturb interactions with other molecules or other parts of the protein. The mutated residue is located in a domain that is important for binding of other molecules and in contact with residues in a domain that is also important for binding. The mutation might interrupt the interaction between these two domains and as such affect the function of the protein.

**Figure 13:**
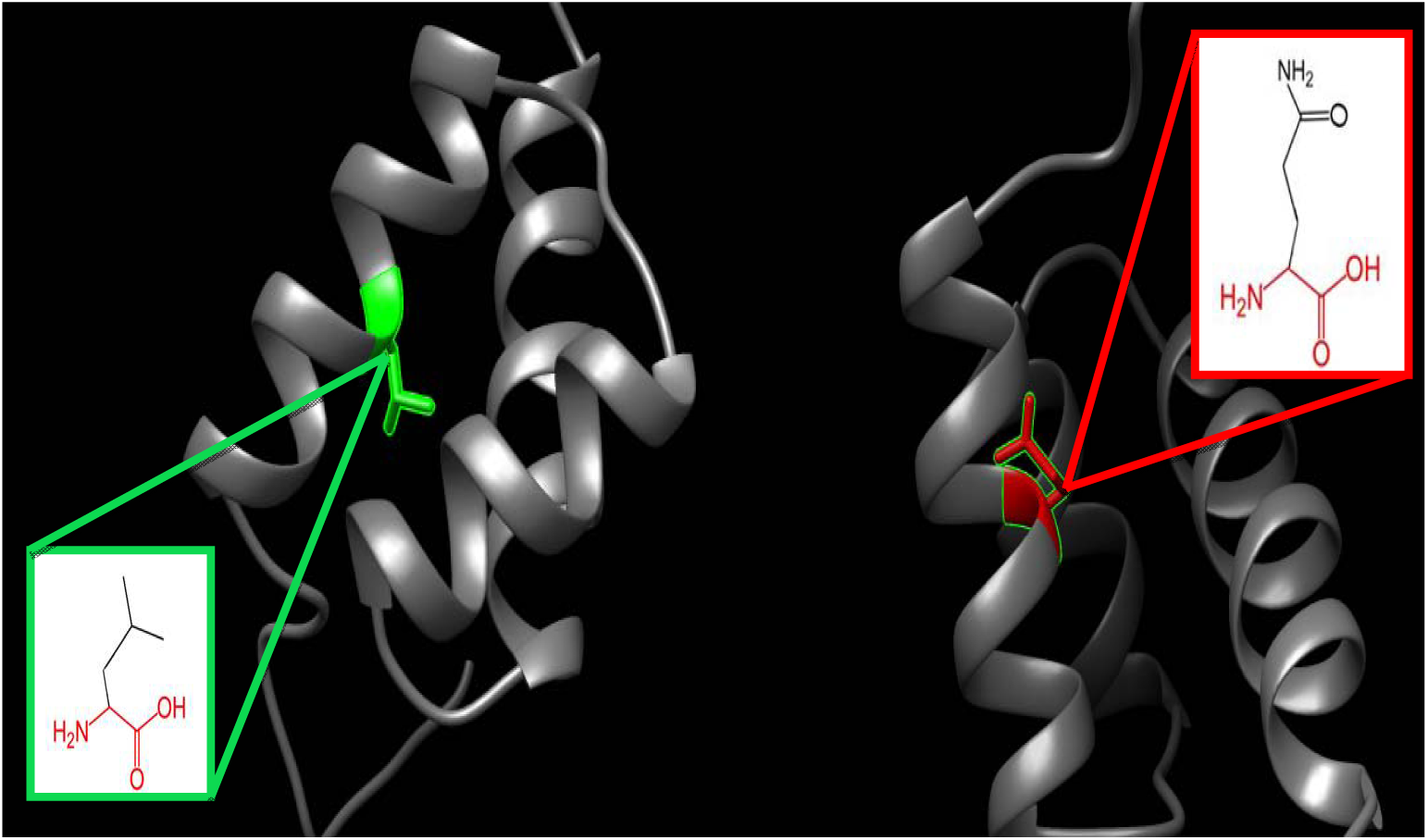
(L343Q): the amino acid Leucine changes to Glutamine at position 343. This visualization was done using Chimera software coupled with the 2D structures (in boxes) from Project Hope report of ARX gene mutations. The green color indicates the wild amino acid, and the red color indicates the mutant one, while the gray color indicates the rest of ARX protein structure.

In **(Figure 14)**: (T333N): the amino acid Threonine changes to Asparagine at position 333; the residue has DNA interactions or is located in a DNA binding region. Threonine was buried in the core of the protein, therefore the mutation will cause loss of hydrophobic interactions in this core. The difference in the hydrophobicity and size between the two amino acids will negatively affect the interactions with associated molecules.

**Figure 14:**
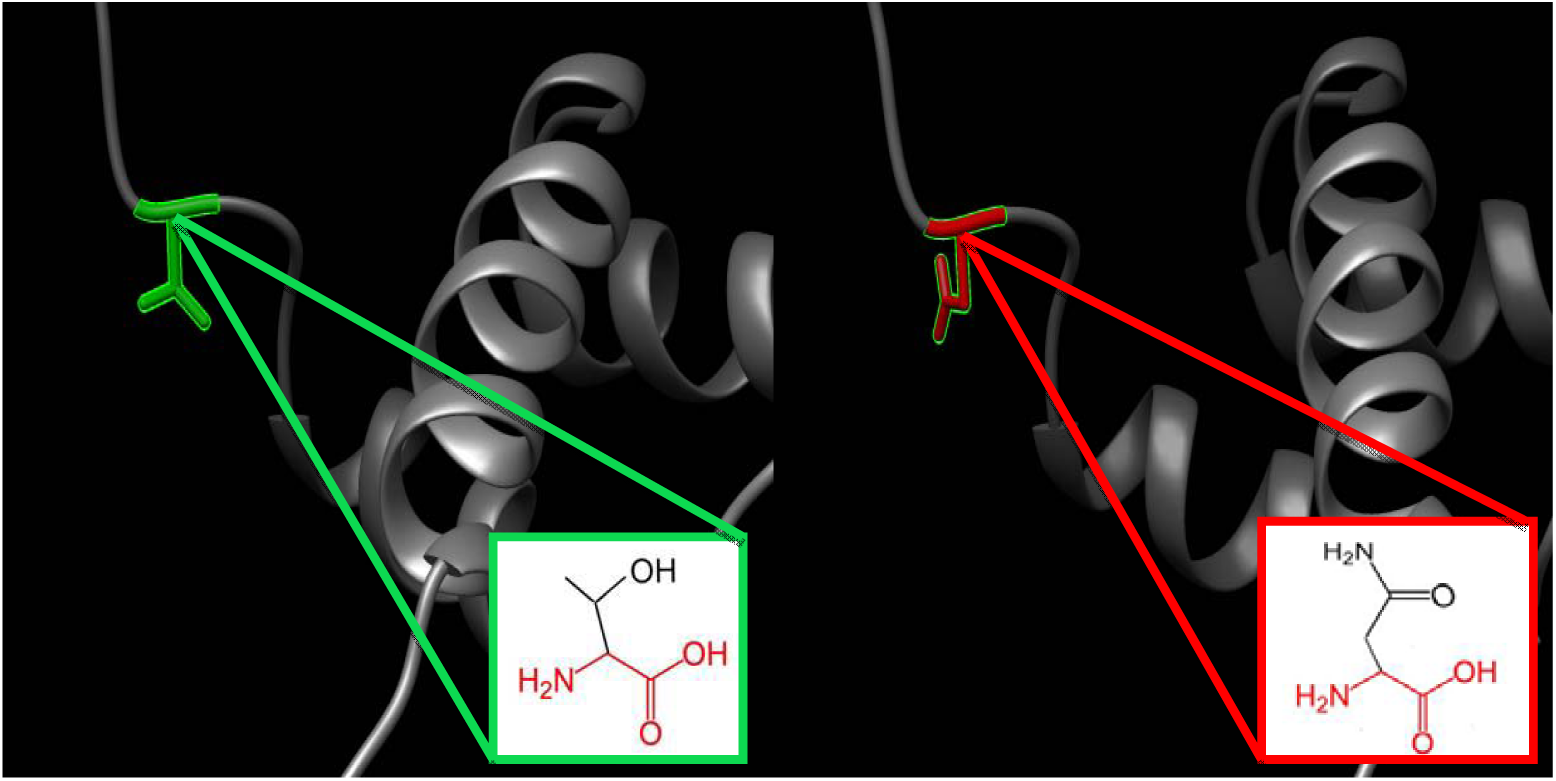
(T333N): the amino acid Threonine changes to Asparagine at position 333. This visualization was done using Chimera software coupled with the 2D structures (in boxes) from Project Hope report of ARX gene mutations. The green color indicates the wild amino acid, and the red color indicates the mutant one, while the gray color indicates the rest of ARX protein structure.

In **(Figure 15)**: (T333S): the amino acid Threonine changes to Serine at position 333; the wild-type and mutant amino acids differ in size. Serine is smaller than the Threonine this will cause an empty space in the core of the protein and defiantly affect its function.

**Figure 15:**
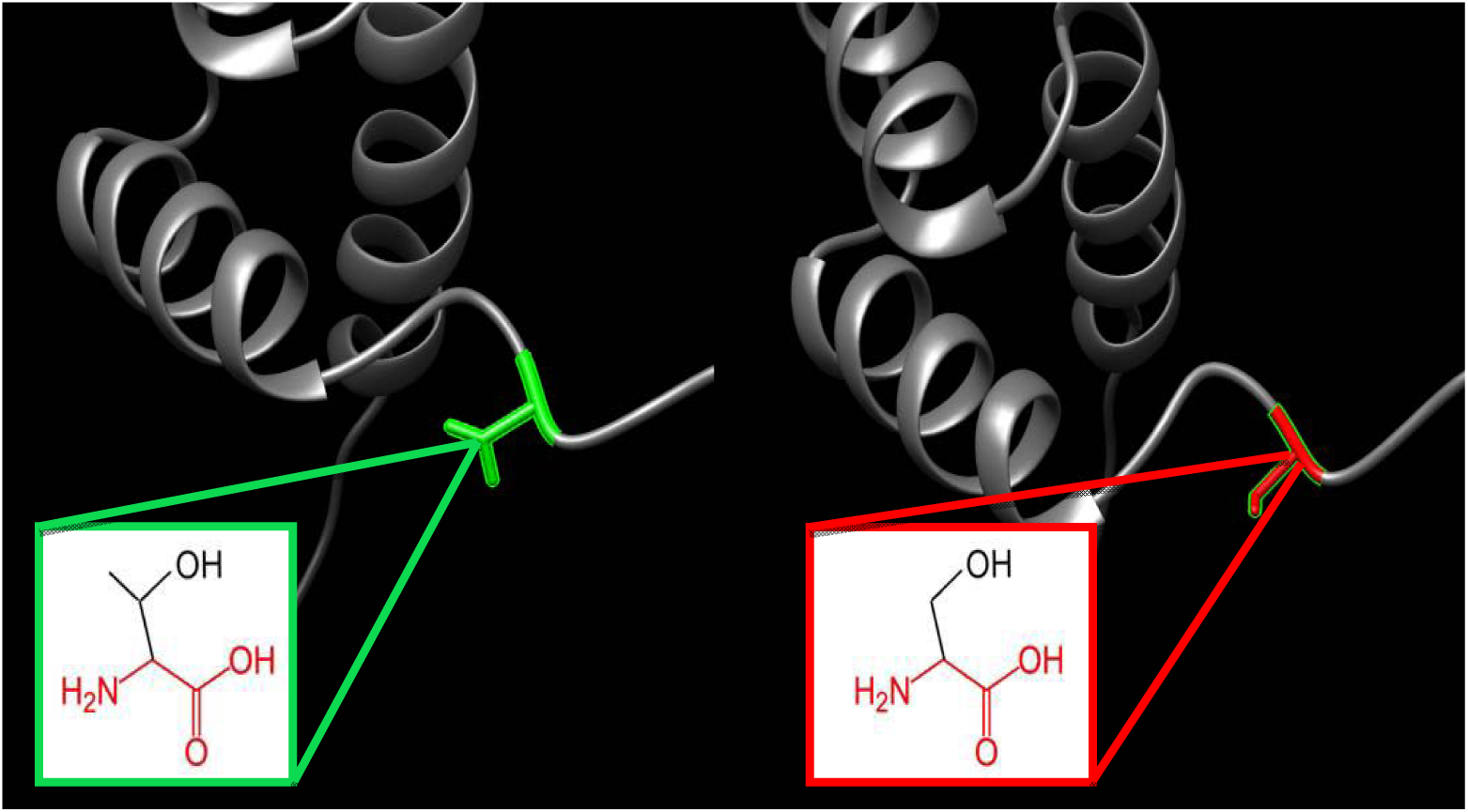
(T333S): the amino acid Threonine changes to Serine at position 333. This visualization was done using Chimera software coupled with the 2D structures (in boxes) from Project Hope report of ARX gene mutations. The green color indicates the wild amino acid, and the red color indicates the mutant one, while the gray color indicates the rest of ARX protein structure.

In **(Figure 16)**: (R332H): the amino acid Arginine changes to Histidine at position 332; Histidine residue is smaller than the wild-type residue. The charge of the wild-type residue (positive) is lost by this mutation. This loss of charge in addition to the smaller size of Arginine will lead to a possible loss of external interactions.

**Figure 16:**
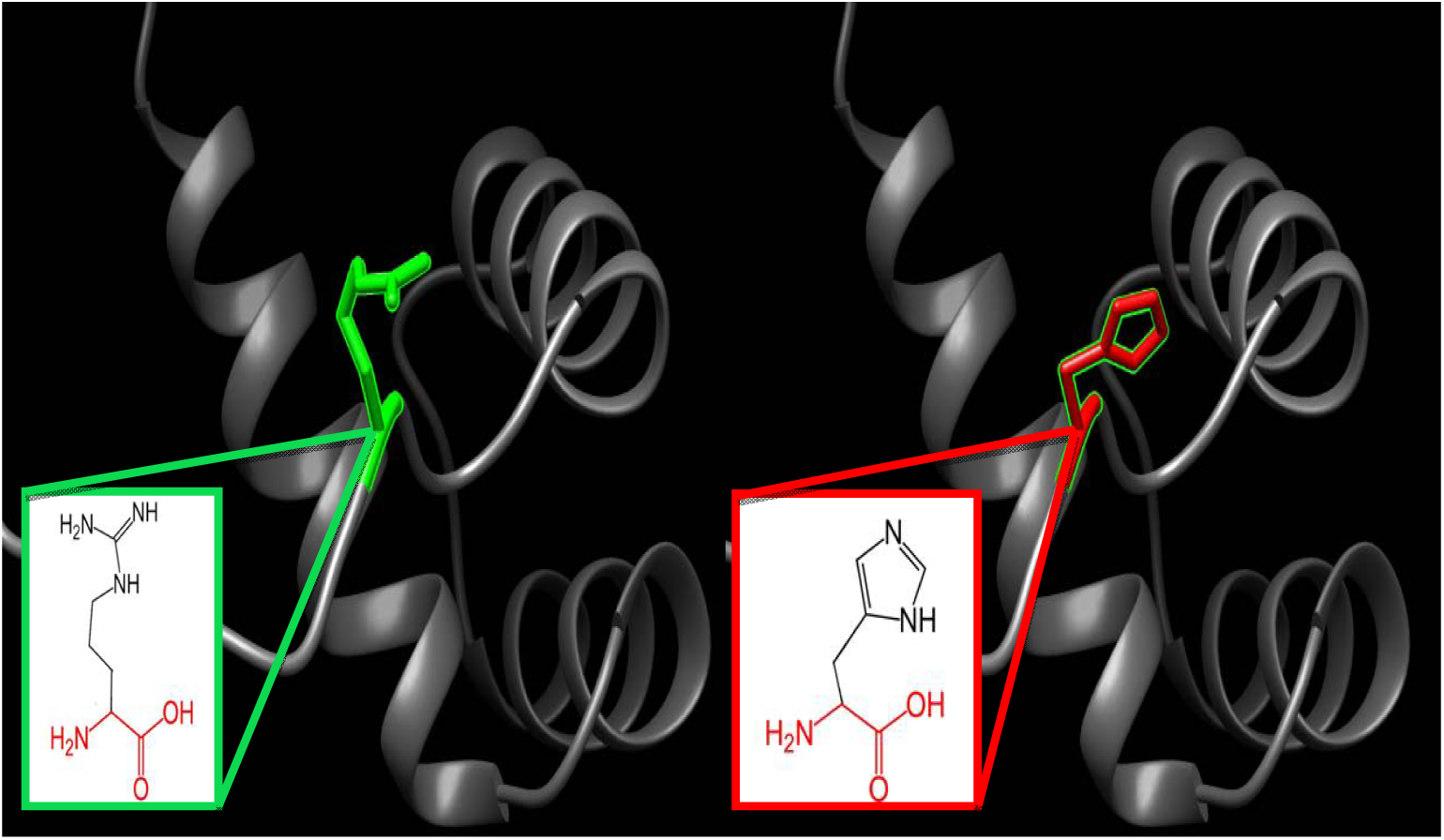
(R332H): the amino acid Arginine changes to Histidine at position 332. This visualization was done using Chimera software coupled with the 2D structures (in boxes) from Project Hope report of ARX gene mutations. The green color indicates the wild amino acid, and the red color indicates the mutant one, while the gray color indicates the rest of ARX protein structure.

In **(Figure 17)**: (R330H): the amino acid Arginine changes to Histidine at position 330; There is a difference in charge between the wild-type (positive) and mutant amino acid (negative); This can cause loss of interactions with other molecules. The mutant residue Histidine is smaller than the wild-type residue; this will negatively affect the external interactions.

**Figure 17:**
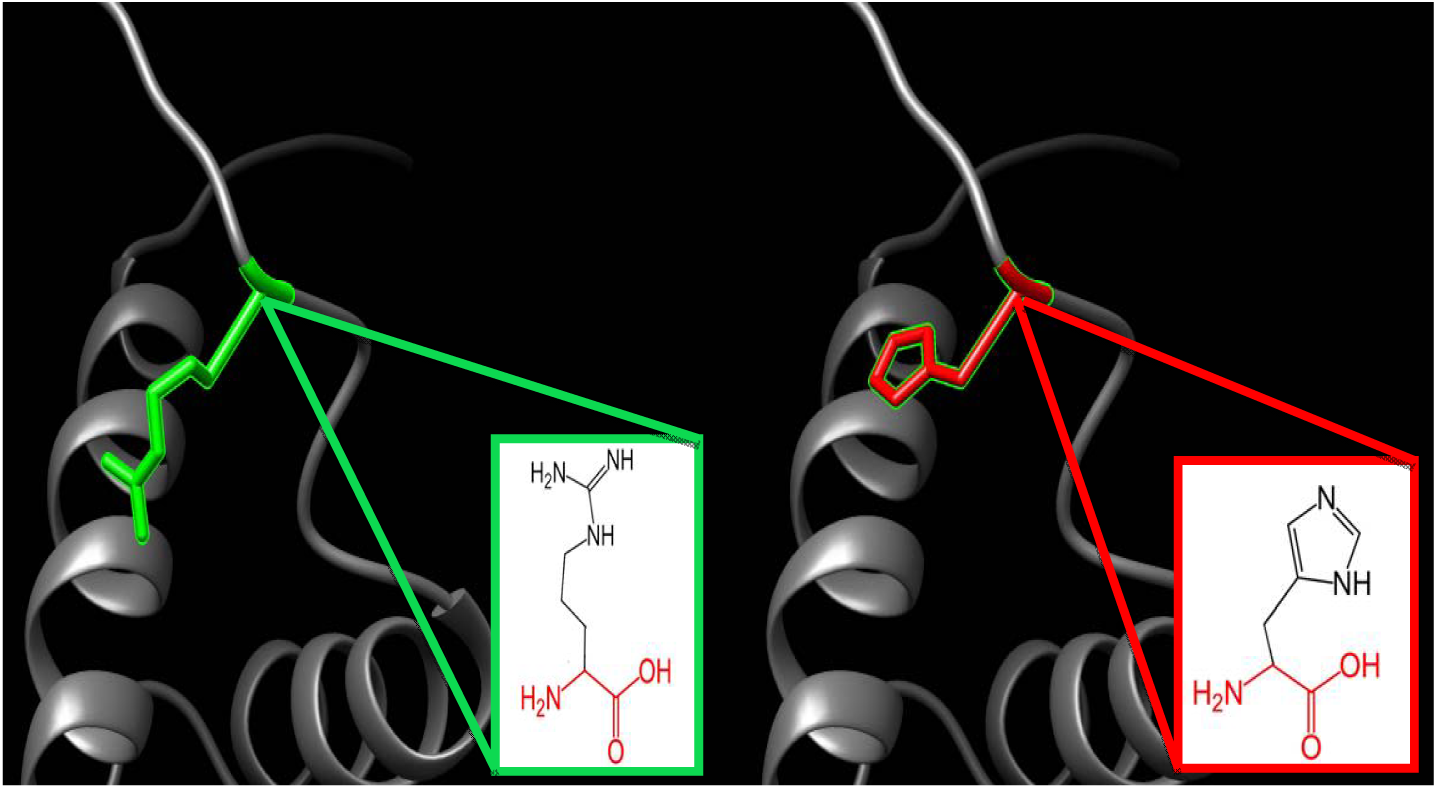
(R330H): the amino acid Arginine changes to Histidine at position 330. This visualization was done using Chimera software coupled with the 2D structures (in boxes) from Project Hope report of ARX gene mutations. The green color indicates the wild amino acid, and the red color indicates the mutant one, while the gray color indicates the rest of ARX protein structure.

As we previously briefed, *ARX* mutations it may cause neuroblastoma; so, to identify the most probable mutations, Mutation3D was used. The following SNPs (R330H, T333N, and T333S) are clustered mutation(colored red), which are commonly associated with human cancers [43]; while these SNPs (L343Q, V373D and R380L) are covered mutations (colored blue) **(Figure 18)**.

**Figure 18:**
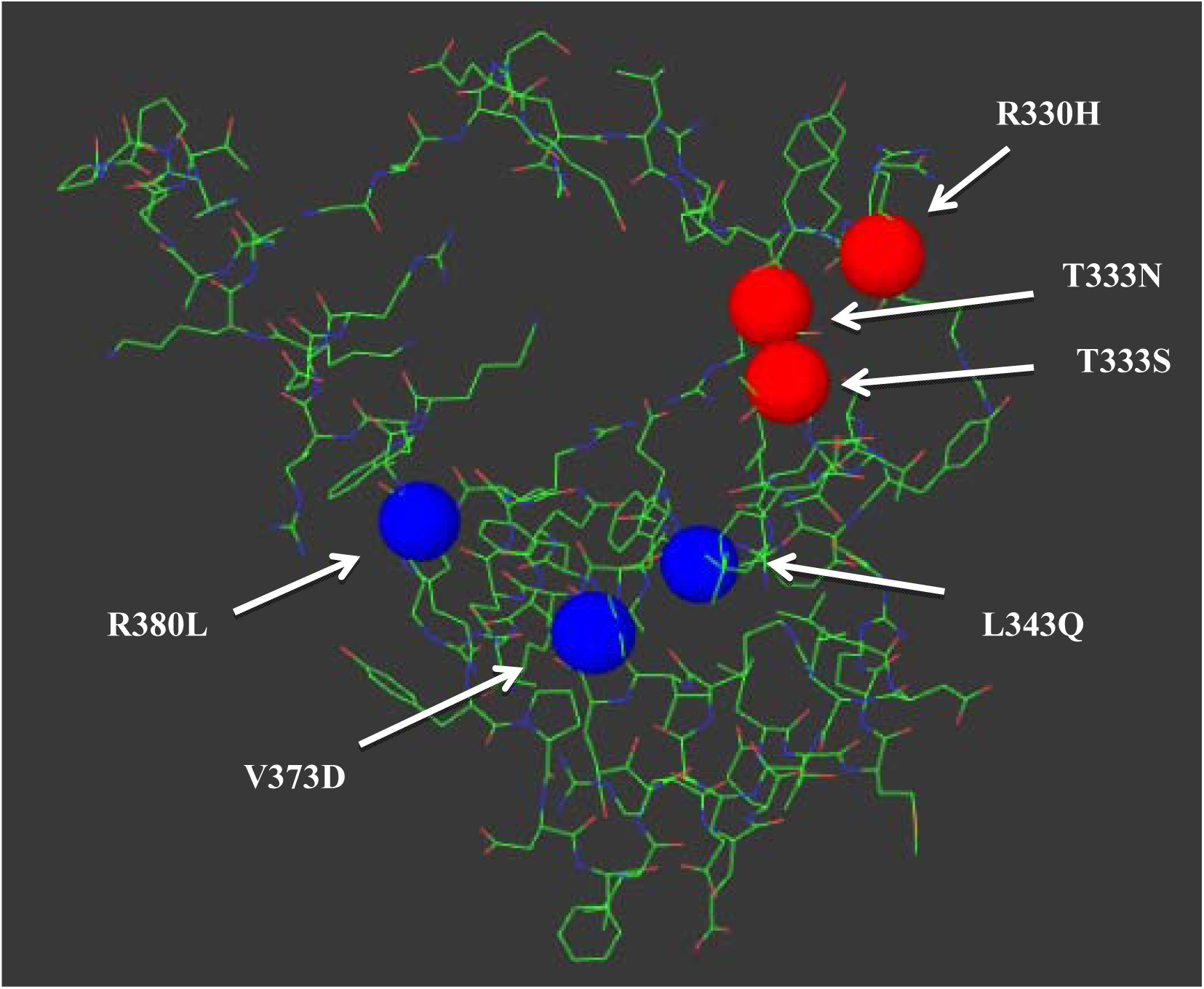
Structural simulations for mutant residues in *ARX* protein predicted by Mutation3D.

We also used ClinVar to compare our results that had been found by an in silico approach with the clinical one; in L535Q, SNP was found to be deleterious, and our result agree with this result [44]; In the second SNP (T333N), the associated clinical studies in ClinVar shows that our result matches with the reported record which is a pathogenic variant [45]; In the third SNP (R332H), the associated clinical study in ClinVar shows that our result matches with the reported record which is a pathogenic variant [7]; while the rest of the most deleterious SNPs (R528S, G461D, R380L, V374D, L343Q, R330H, G34R and L33P), we did not find any associated clinical studies.

This study is the first computational analysis of *ARX* gene while all other previous studies were in other approaches [46-48]. The limitation of this study is that it focuses on coding region of *ARX* gene using several numbers of computational analysis tools. Although EIEE1 is frequently triggered by mutation in *ARX* gene [6]; yet there are number of genes responsible for EIEE1 [46, 49]; another limitation is that the 3D structure of the ARX protein model was created by using RaptorX; it could not create the 3D structure of all amino acid positions. Therefore, the model was done from position 325 to 390, due to the lack of information.

To conclude, *in silico* analysis of the functional and structural consequences of SNPs in human ARX gene revealed 11 mutations that may responsible for causing EIEE1. It is obvious to realize that bioinformatics approach remains a cost effective way to make a rapid analysis regarding the expected effect of variants; yet, the more factors that are taken into account, the more accurate the prediction will be. Finally, clinical wet lab studies are recommended to support these findings

## 5. Conclusion

Extensive *in silico* analysis of the functional and structural consequences of SNPs in human ARX gene revealed 11 mutations (L535Q, R528S, R380L, V374D, L343Q, T333N, T333S, R332H, R330H, G34R and L33P) that may cause EIEE1 disease. These can be used as diagnostic markers for EIEE1 or as a possible drug targets.

## Data Availability

All data underlying the results are available as part of the article, and no additional source data were required.

## Acknowledgment

The authors acknowledge the Deanship of Scientific Research at University of Bahri for the supportive cooperation.

## Authors’ contributions

MIM: Illustration and drafted the manuscript; NSM & AHA: collected, analyzed and interpreted the data; AMM: designed and revised (review and editing) the work. All authors have read and approved the final paper.

